# Axial regionalization in *Tiktaalik roseae* and the origin of quadrupedal locomotion

**DOI:** 10.1101/2023.01.11.523301

**Authors:** T.A. Stewart, J.B. Lemberg, E.J. Hillan, I. Magallanes, E.B. Daeschler, N.H. Shubin

## Abstract

The origin of quadrupedal locomotion in tetrapods entailed the evolution of a regionalized axial skeleton with sacral ribs. These ribs provide linkage between the pelvis and vertebral column and contribute to body support and propulsion by the hind limb. The closest relatives of limbed vertebrates are not known to possess such a connection and, therefore, have typically been described as primarily supporting their bodies against the substrate with pectoral fins. However, data on the axial skeletons of stem tetrapods are sparce, with key features of specimens potentially concealed by matrix. Here we provide micro-computed tomography data of the axial skeleton of *Tiktaalik roseae* and show that its vertebrae and ribs are regionalized along the craniocaudal axis, including expanded and ventrally curved ribs in the sacral region. The sacral ribs would have contacted the expanded iliac blade of the pelvis in a soft tissue connection. No atlas-axis complex is observed, however the basioccipital-exoccipital complex is deconsolidated from the rest of the neurocranium, suggesting increased mobility at occipital-vertebral junction. Thus, axial regionalization that allowed for innovations in head mobility, body support and buttressing the pelvic fin evolved prior to the origin of limbs.

---

The earliest limbed vertebrates are characterized by a regionalized axial skeleton with cervical, thoracic, sacral, and caudal domains in the vertebral column and ribs^1–4^. These modules correspond to locomotor specializations, including providing support for load-bearing hind limbs^5–7^. *Acanthostega* and *Ichthyostega* have specialized ribs that connected to the ilium, either in a soft-tissue or bony articulation, providing mechanical linkage between the axial column and the pelvic girdle^1,2,4^. The structures that make this connection possible are not present in tetrapodomorph outgroups where the complete axial column has been described^8–10^. For example, the tristichopterid *Eusthenopteron* has vertebrae and ribs that are short and generally similar across their cranio-caudal distribution, lacks a sacral rib, and has a pelvis that is small as compared to the pectoral girdle^8^. Moreover, unlike *Acanthostega* and *Ichthyostega, Eusthenopteron* possessed a bony linkage between the shoulder girdle and cranium that would have limited head mobility^1,2,4,8^.

Little is known of the axial columns of the closest relatives of limbed vertebrates. The vertebrae of *Panderichthys* are described from a brief series that evince no indication of regionalization^11^. The vertebrae of *Elpistostege* are known from a series of approximately 16 that, likewise, show no heterogeneity in their length or shape^12^. The axial skeleton of *Tiktaalik* has been largely obscured by matrix. The rostral ribs are broad and laterally expanded as compared to early tetrapodomorph conditions, and vertebral column has not been described^13^. However, the pelvis and pelvic fin of *Tiktaalik* are nearly the size of the pectoral appendage^14^, differentiating its overall proportions from less crownward taxa, like *Panderichthys^11,15^*. The size and depth of the acetabulum, the general robusticity of the pubis, and the dorsally expanded iliac blades of *Tiktaalik* are further similarities shared with digited forms that are notably absent in other finned tetrapodomorphs^14^.

Here, we present high-resolution micro-computed tomography scans of the type specimen of *Tiktaalik*, NUFV 108, that expose for the first time the vertebral skeleton and posterior ribs of *Tiktaalik* (Fig. 1, Movies S1,2). These data, and the new reconstruction they allow, reveal unexpected intermediate conditions as well as apomorphies that provide new insight on changes involved with the origin of limbed vertebrates and the functional context in which they arose.

**Fig. 1.**
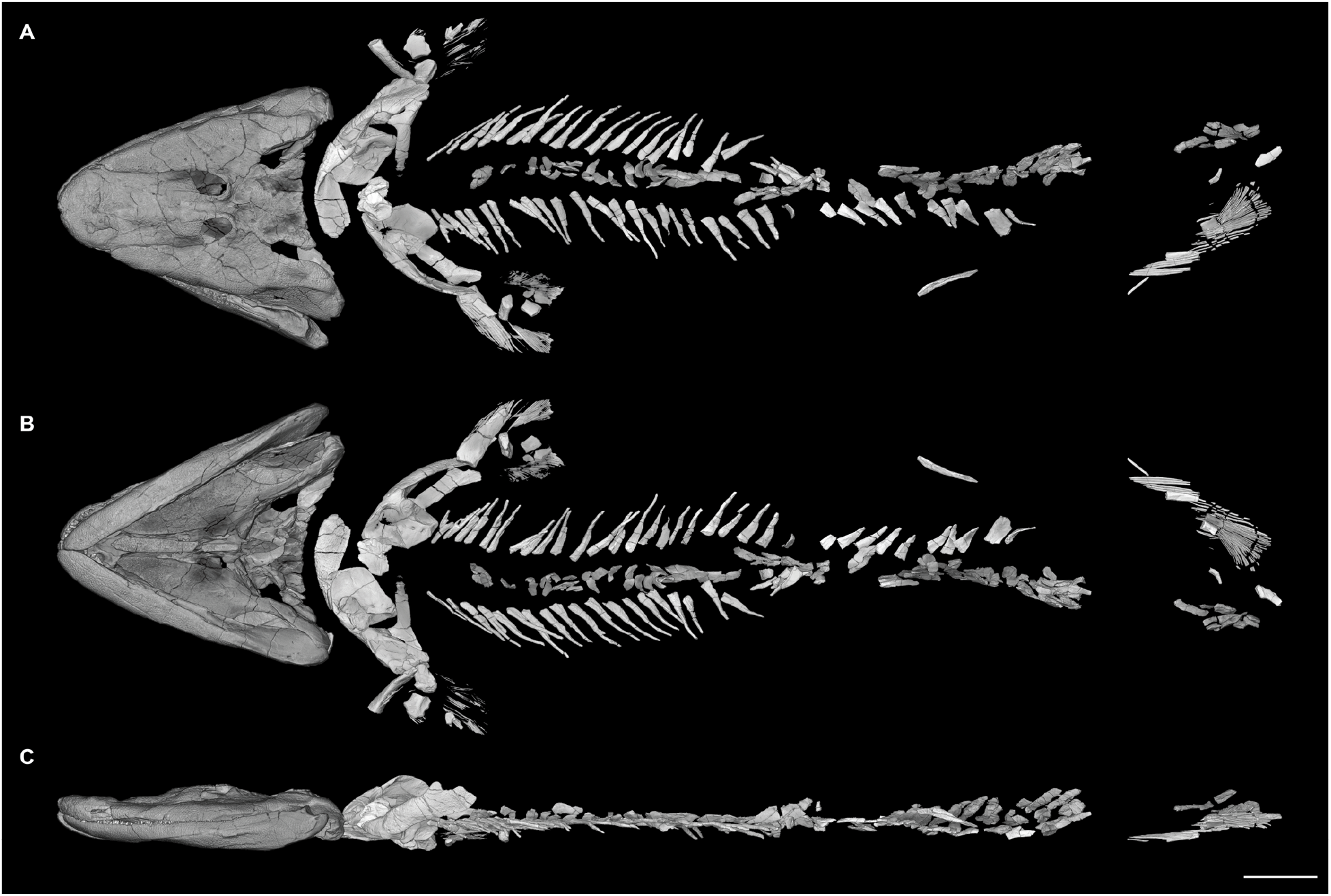
Volumetric rendering of μCT scans of *Tiktaalik roseae*. Specimen NUFV 108 in (**A**) dorsal, (**B**) ventral, and (**C**) left lateral perspectives. μCT data reveal new detail on the ribs, vertebrae, and pelvic fin. The head, which was mechanically prepared and scanned separately^27^, is positioned slightly anterior to its preserved position. Scale bar, 5 cm.

## Results

### Vertebrae

The vertebrae of *Tiktaalik* are rhachitomous and surround an unconstricted notochord that was persistent into adulthood (Fig. 2). In specimen NUFV 108, elements of 40 vertebrae are preserved. These include ossified intercentra and neural arches, while pleurocentra are not identified. The size, shape, and spacing of intercentra and neural arches of *Tiktaalik* are similar to *Eusthenopteron^1^*, suggesting that pleurocentra might have been present but, because of their small size, are difficult to identify within the field of preserved scales. However, their absence in *Tiktaalik* cannot be excluded; pleurocentra have likewise not been identified in *Panderichthys^11^* or *Elpistostege*^12^, suggesting a possible ‘reverse’ rhachitomous pattern in elpistostegalians which is known in *Acanthostega, Ichthyostega,* and *Pederpes^3^*.

**Fig. 2.**
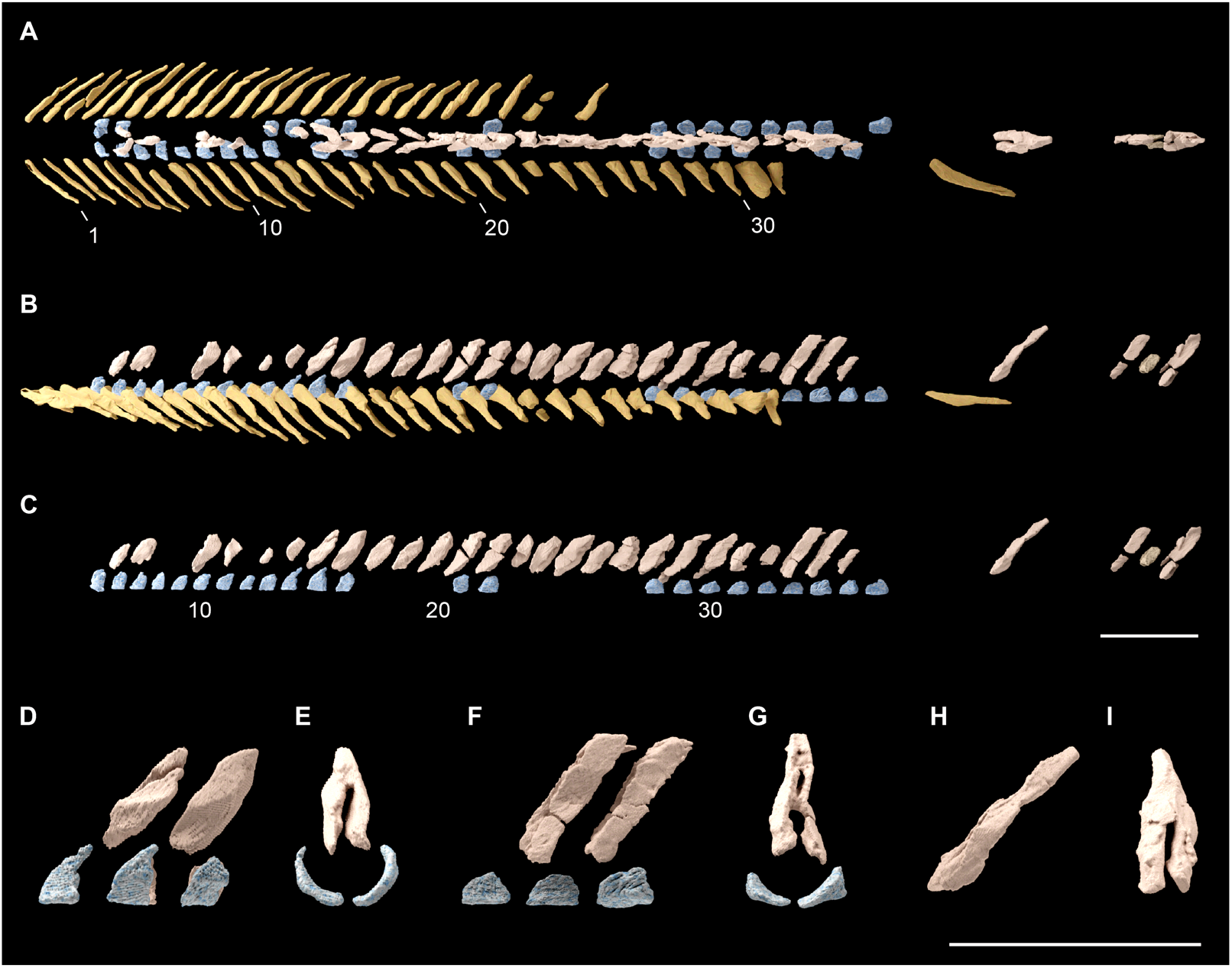
Vertebrae and ribs of *Tiktaalik roseae*. Vertebrae and ribs in (**A**) dorsal and (**B**) lateral perspective. (**C**) Intercentra and neural arches in lateral perspective. (**D,E**) Intercentra and neural arches beginning at position 14 in left lateral and anterior perspective (**F,G**) Intercentra and neural arches beginning at position 32 in left lateral and anterior perspective. (**H,I**) Neural arch from the caudal region in left lateral and anterior perspective. Ribs are depicted in yellow, neural arches in tan, and intercentra in blue. Scale bars, 5 cm.

Vertebrae are not preserved in association with the four most anterior ribs (Fig. 2 A,B). It is possible that these vertebrae were present but are not preserved in specimen NUFV 108. However, a similar condition is observed *Ichthyostega*, where preserved intercentra and neural arches also initiate at rib number five^3^. This well-defined gap in multiple taxa suggests that these vertebrae in the cervical domain were cartilaginous into adult stages in *Tiktaalik* and *Ichthyostega* and that the observed pattern is not an artifact of preservation or variation across ontogeny.

Intercentra are paired and have minor graded differences in their morphology across the series (Fig. 2 C). Proceeding caudally, intercentra become longer in the rostro-caudal direction, shorter dorsoventrally, and bear a larger articular facet for the ribs (Fig. 2 D-G). Similar rostro-caudal variation is observed in the presacral intercentra of *Eusthenopteron^8^*. *Tiktaalik* is distinguished from closely related taxa in having paired intercentra along the full series. In *Eusthenopteron,* the anterior five intercentra and the intercentra above the pelvis, at approximately position 32, are bilaterally fused^8^; *Acanthostega* has fused atlantal and sacral intercentra^1^; and in *Ichthyostega* most intercentra are fused, with only the anterior-most ones being paired^3^.

Neural arches are inclined posteriorly and vary craniocaudally in their morphology. Frequently, they are laterally compressed in preservation, with the left and right halves occasionally separating, as has been described in *Eusthenopteron*^8^, *Panderichthys*^11^, *Elpistostege*^12^, and *Acanthostega^1^*. Zygapophyses are not observed, unlike the condition of limbed *vertebrates^2,3,16,17^*. Cranially, neural arches have a simple saddle shape (Fig. 2 D,E). The rostral 30 arches show subtle variation in their geometry, with more caudal neural arches having slightly more vertical inclination relative to the notochord. By position 32, the neural arch pattern shifts abruptly, and neural arches extend further dorsally and have a dorsal foramen. Neural arch 31 is broken dorsally, and so it is unclear whether the transition in neural arch morphology occurs at position 31 or 32. Regardless, this shift in morphology is inferred to mark the trunk-tail boundary, a change also observed in the ribs as described below. Further caudally, four vertebrae are preserved. One of these is substantially more robust than all others (Fig. 2 F,G. Movie S1), similar to neural arches preserved in the caudal domain of *Acanthostega^1^*.

### Ribs

Specimen NUFV 108 was physically prepared in 2004 and 2005 to expose rostral ribs^13^. μCT imaging reveals additional ribs preserved beyond those previously identified, making for a total of 56, including an uninterrupted series of 32 on the left side. Across the series, ribs have a curved articular head that would have contacted the pleurapophyses of the intercentra. Ribs bear a flange posteriorly on their proximal portion that varies in its mediolateral span across the series, and they lack imbricating uncinate processes (Fig. 2 A,B). The rostral-most ribs extend straight to a tapered, narrow tip. More caudally, at approximately rib number 5, the ribs become longer and have a gentle ventral curvature. At approximately rib number 20, the ribs shorten in their mediolateral span and have a broader base, gaining a more triangular shape. Ribs 31 and 32 are markedly distinct in their morphology from others in the series. Rib 31 is broad in dorsal perspective and has unfinished distal surface that is rounded, while rib 32 shows substantial ventral curvature as compared more cranial ribs (Fig. 2 A,B). An isolated post-sacral rib is preserved to the left of the other axial elements (Fig. 1 A,B, Movie S2). Its morphology, narrow, slightly recurved and posteriorly directed, is similar to the post-sacral ribs of *Acanthostega^1^* and *Ichthyostega^2^*. No evidence of sternal structures is found.

### Reconstruction of the pelvic region

The morphology of the pelvis of *Tiktaalik* was described previously^14^, but, importantly, its position and relation to the axial column has remained unknown. The right pelvis of specimen NUFV 108 was preserved adjacent to the axial column, not in articulation^14^. Abrupt transitions in the morphology of the vertebrae and ribs at position 31 and 32 in *Tiktaalik* denote the trunk-to-tail transition and likely position of the pelvic girdle. In both *Eusthenopteron* and *Acanthostega*, transitions in vertebral and rib anatomy at this general position denote the trunk-to-tail transition and pelvic position. In *Eusthenopteron*, ribs are only present rostral to vertebrae 30, between vertebrae 30 and 32 the haemal arches enclosed the haemal canal and the become intercentra fuse bilaterally, and the pelvis is approximately ventral to vertebrae 32^8^. In *Acanthostega*, vertebrae 31 differs from those immediately rostral in having fused intercentra and bearing a distinctive and elongate rib with a ventral expansion that would have allowed for attachment to the girdle, likely *via* soft tissue^1^.

New data on the on vertebrae and ribs of *Tiktaalik* allow for assessment of pelvic orientation and position (Fig. 3, Movie S3). The dorsal extent of the iliac blade of the pelvis would have approached ribs 31 and 32 of the series. While there is no articular facet on the internal surface of the girdle for a sacral rib^14^, the proximity inferred from their anatomy, along with a comparison to other taxa, suggest a soft-tissue linkage. In *Acanthostega*, there is likewise no distinct articular facet or marked perimeter for the attachment of the sacral rib in the ilium^1^. Nevertheless, a soft-tissue connection between rib and ilium has been inferred based on rib morphology and the size and position of the two elements^1^. Similar patterns of connectivity have been proposed in other early tetrapods, such as *Eryops*^1,18^. A bony articulation between ribs and pelvis is only definitively present in more crownward tetrapods such as *Whatcheeria^1,18,19^*. The positioning of the pelvis of *Tiktaalik* suggested by the shape of the pubis and width of the body (Fig. S1, Supplementary Text) would entail a more posteroventral-facing acetabulum than previously proposed^14^, more similar to the orientation of the pelvic fins of *Eusthenopteron*^8^ than the laterally positioned limbs of Devonian limbed vertebrates^1,2,20^.

**Fig. 3.**
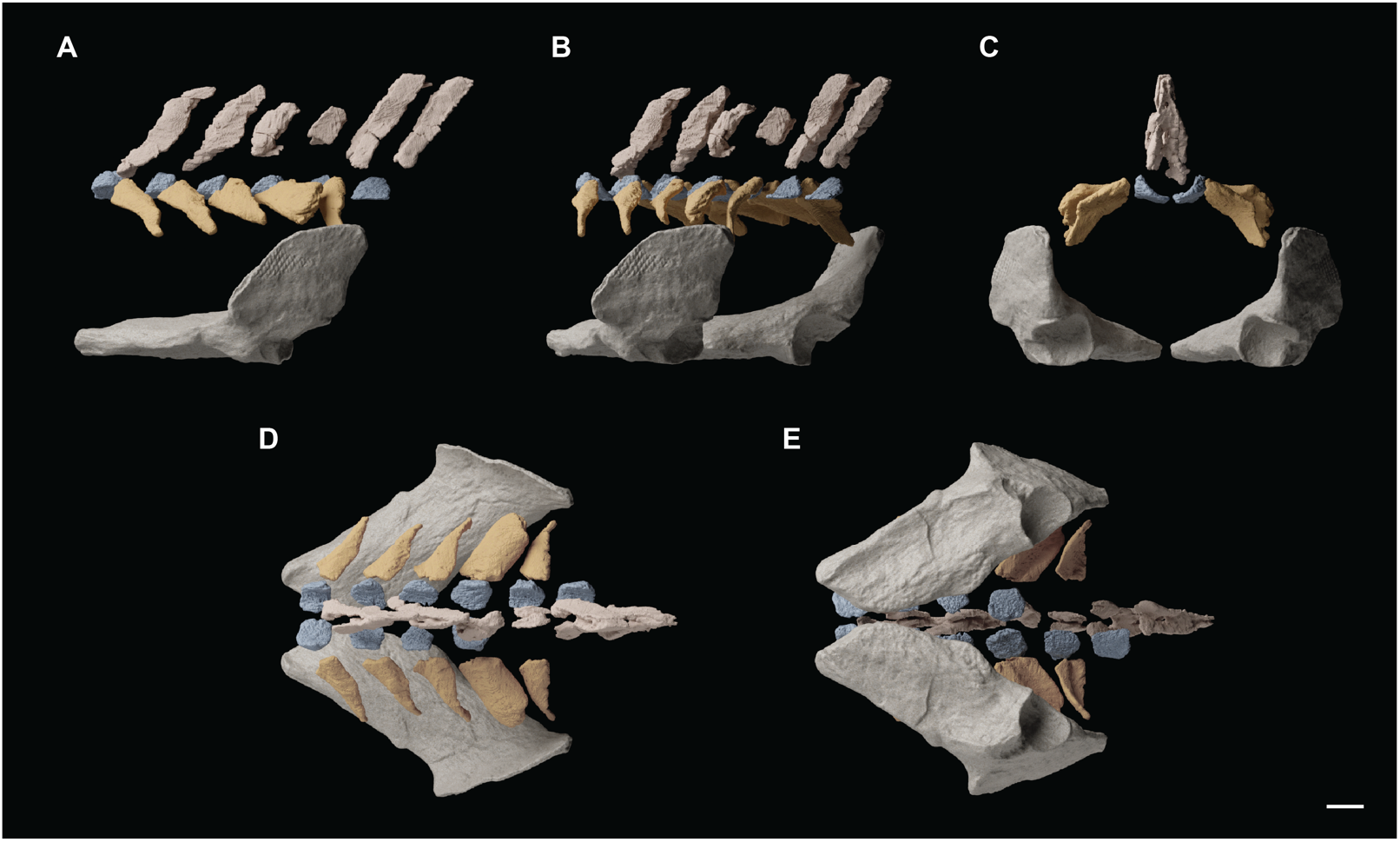
Reconstructed sacral domain and pelvic girdle of *Tiktaalik roseae*. Reconstruction of the axial column and pelvis in (**A**) left lateral, (**B**) posterior-oblique, (**C**) posterior, (**D**) dorsal, (**E**) ventral perspectives. Ribs and pelvic girdle have been mirrored to produce the reconstruction. Ribs 31 and 32 show modified shape as compared to the more anterior elements and are inferred to have supported the pelvic girdle by a soft-tissue connection. Scale bar, 1 cm.

### Pelvic fin

Mechanical preparation of specimen NUFV 108 in 2005-2006 exposed parts of the pelvic fin^14^. μCT data reveal new details, including the full extent of the pelvic fin web and additional endoskeletal elements (Fig. 1, Fig. 4 A). Pelvic fin rays are unbranching and unsegmented.

**Fig. 4.**
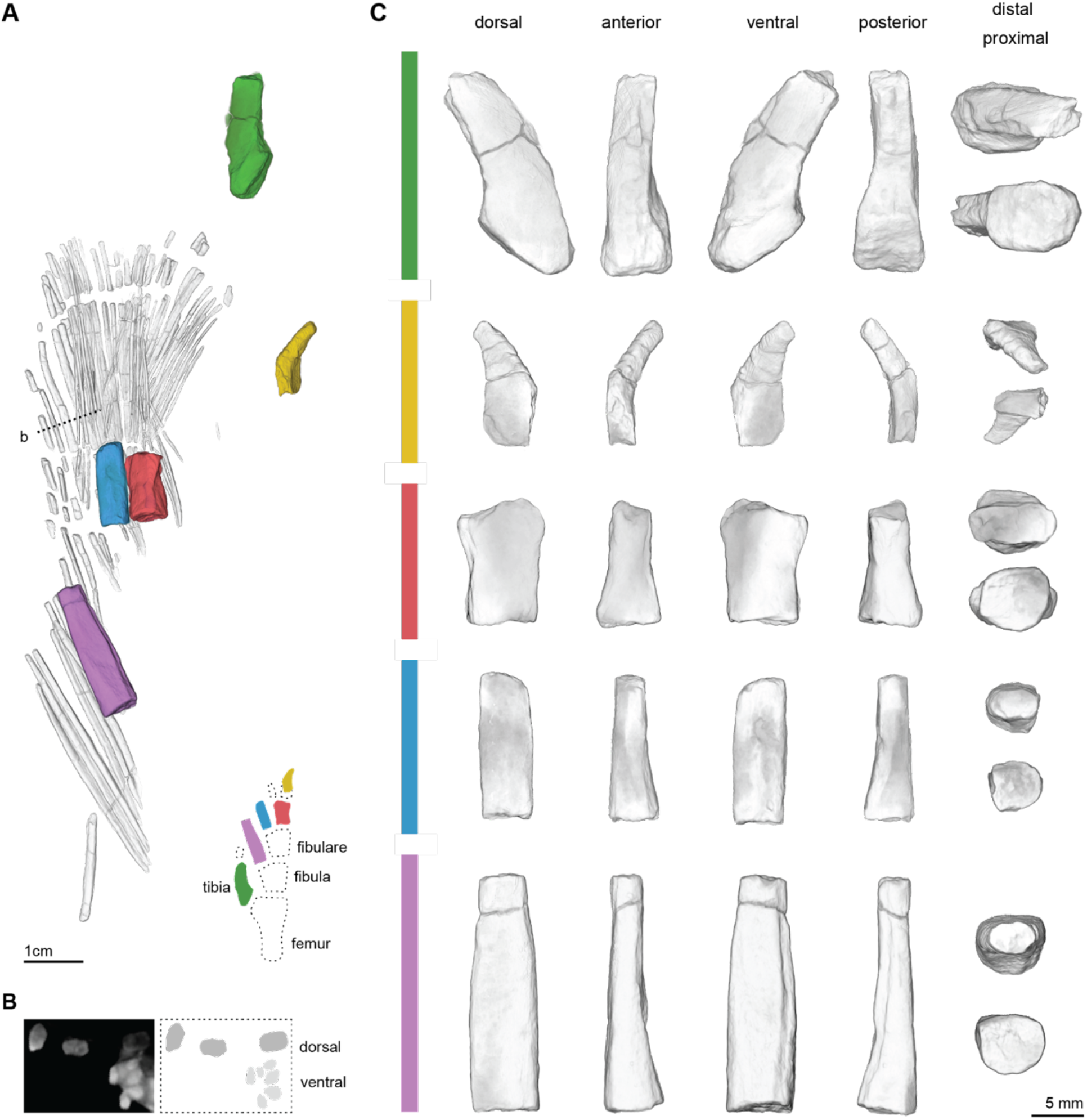
Pelvic fin of *Tiktaalik roseae*. (**A**) Volumetric rendering of μCT data of the left pelvic fin of NUFV 108 and a reconstruction of the fin in ventral perspective. (**B**) Hemitrichia show dorso-ventral asymmetry. The digital cross section, left, and illustration, right, were taken at the position of the dashed line labeled ‘b’ in panel A. The cross section is oriented orthogonal to the plane of the fin web. (**C**) Endoskeletal elements of the pelvic fin in various orientations.

Similar to the pectoral fins of tetrapodomorphs, the pelvic fin rays are more robust on the leading edge and more gracile on the posterior side^21^. Hemitrichia have accentuated asymmetry. Dorsal hemitrichia are larger in cross section than ventral hemitrichia, as in the pectoral fin of *Tiktaalik*^21^ (Fig. 4 B). Two new pelvic endoskeletal elements are identified (Fig. 4 A,C). One, inferred to be a tibia, has a robust proximal articular surface, and its distal margin appears broken, making it unclear whether a more distal element might have articulated with it. The other element is small with a posteriorly oriented ventral curving process, a feature not previously observed in tetrapodomorph pelvic fins^8–10,15,22^.

### Occipital-vertebral junction

In specimen NUFV 108, the basioccipital-exoccipital complex is preserved apart from the rest of the skull, medial to the pectoral girdles, and it comprises a bilateral pair of elements (Fig. 1, Fig. S2 A-D). Examination of μCT data of specimen NUFV 110^23^ confirms that the basioccipital-exoccipital complex is deconsolidated from the rest of the skull in *Tiktaalik*. (Fig. S1 E-I). The pattern of *Tiktaalik* differs from the general pattern among tetrapodomorphs, where the basioccipital-exoccipital complex is fused both across the midline and to anterior neurocranial elements^24,25^. In the tristichopterid *Mandageria fairfaxi*, the basioccipital-exoccipital complex is also separated from more anterior elements, and this feature has been inferred to allow for increased notochordal flexion at the occipital-vertebral junction^26^. Deconsolidation of skeletal elements at the back of the skull in *Tiktaalik*, therefore, provides further evidence for increased mobility at the head-trunk boundary, which was previously hypothesized based on the absence of an operculum and extrascapular series^13^.

## Discussion

*Tiktaalik* exhibits a unique constellation of primitive and derived characters in the axial skeleton that suggest it had a locomotor capacity intermediate to currently known finned elpistostegalians and limbed vertebrates. These data, and the reconstruction they imply (Fig. 5), allow for new hypotheses on the evolution of axial regionalization and the origin of quadrupedal locomotion in early tetrapods.

**Fig. 5.**
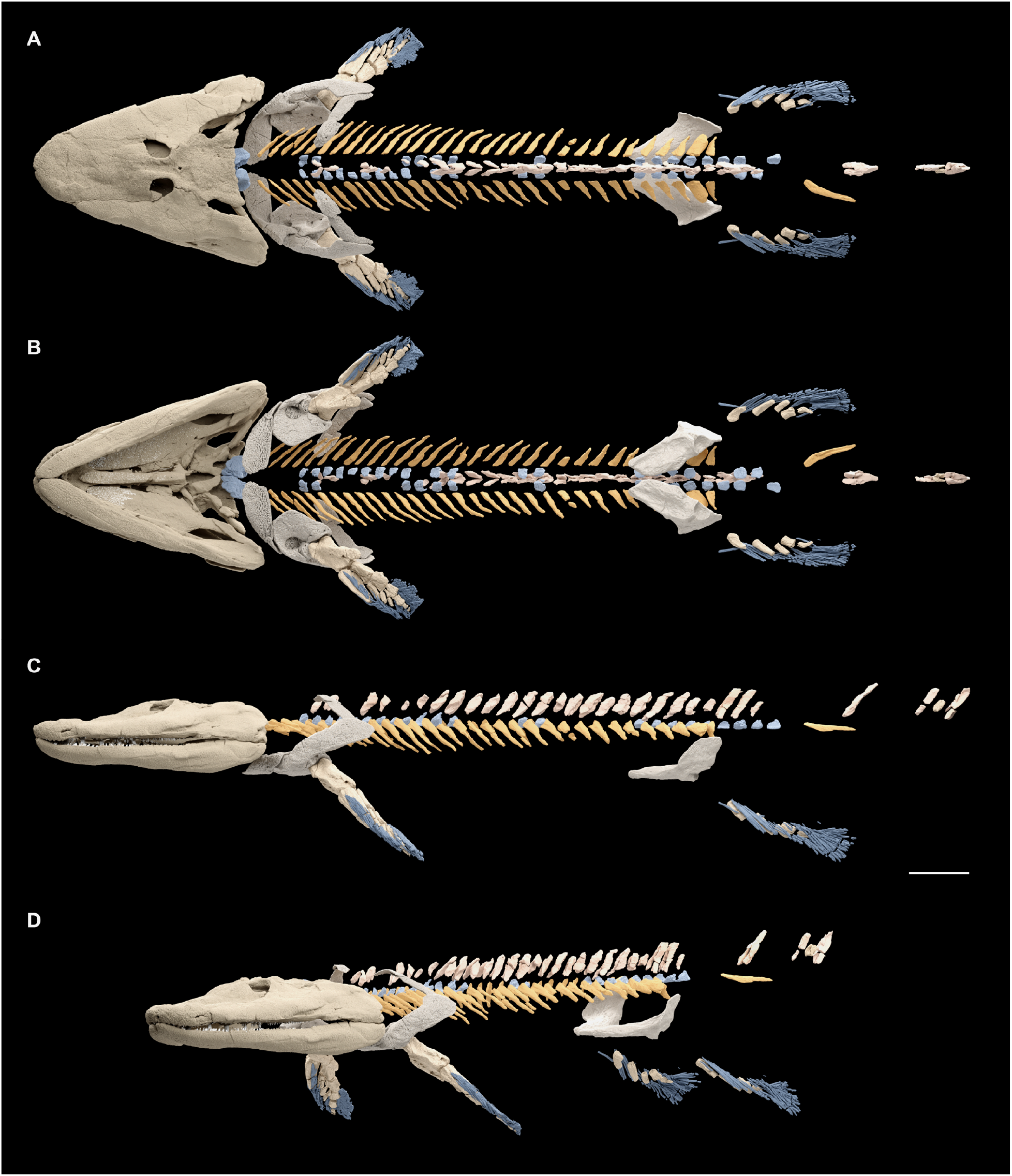
Reconstruction of *Tiktaalik roseae*. Reconstruction in (**A**) dorsal, (**B**) ventral, (**C**) left lateral, and (**D**) oblique views. Cranial materials are repositioned according to Lemberg *et al*^23^ to account for settling during preservation. Select elements that are preserved from only one side of NUFV 108 (*i.e*., pre-sacral ribs, pelvic girdle, and fin) are reflected for symmetry. The pectoral fin is from specimen NUFV 110^21^ and scaled to the length of the right humerus of NUFV 108. Additional skeletal elements are known for *Tiktaalik*, including branchial skeleton^13^ and interclavicle^28^, but have not been rendered here. Scale bar, 5 cm.

The vertebrae of *Tiktaalik* adhere closely to plesiomorphic tetrapodomorph conditions. Most of the preserved vertebrae are from the trunk, and they are similar to the trunk vertebrae of *Eusthenopteron* both in degree of differentiation across the series and in overall construction, except for slight differences in interercentral fusion and the potential lack of pleurocentra^8^. The number of trunk vertebrae in *Tiktaalik* is similar to other tetrapodomorphs; *Eusthenopteron*, *Acanthostega*, and *Ichthyostega* are also characterized by approximately 30 pre-sacral vertebrae^1,2,8^.

In contrast to the vertebral column, the ribs of *Tiktaalik* show numerous derived features that are previously known only from limbed taxa (Fig. 6). As in *Acanthostega*^1^ and *Ichthyostega*^2^, the ribs of *Tiktaalik* extend caudal to the trunk-tail boundary and are regionalized with a sacral module. This is a departure from plesiomorphic tetrapodomorph pattern, seen in *Eusthenopteron*, where ribs do not extend caudal to the trunk-tail boundary and those near to the pelvis are not morphologically differentiated^8^.

**Fig. 6.**
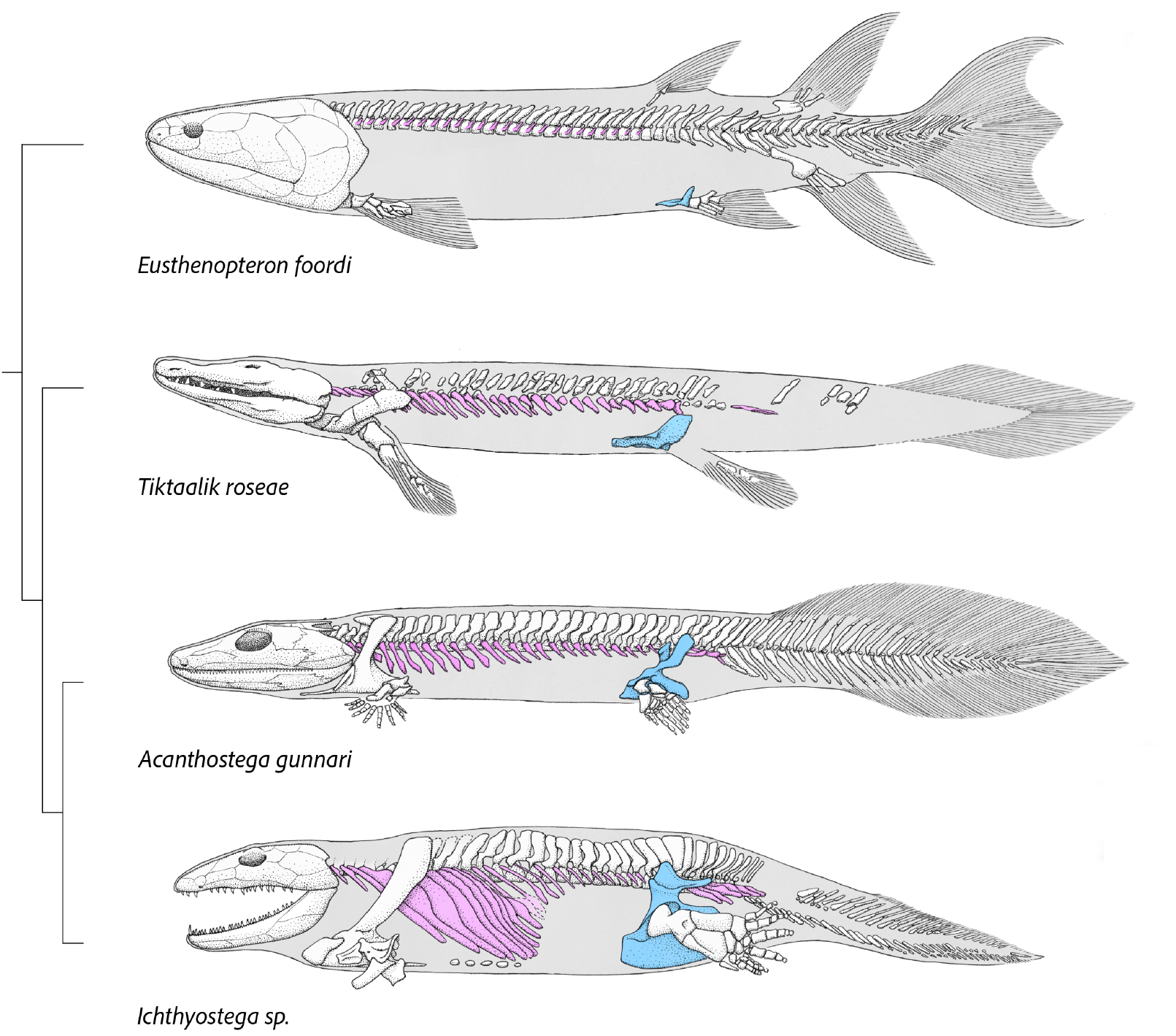
Reconstructions of Devonian tetrapodomorphs. The anatomy of *Tiktaalik roseae* shows that specializations in the axial column for head mobility, body support and pelvic fin buttressing had evolved in elpistostegalians, prior to the origin of limbs. Illustration of *Tiktaalik roseae* based on specimen NUFV 108. Illustrations of other taxa are based on previously published descriptions: *Eusthenopteron foordi*^8,24^, *Acanthostega gunnari*^1,2^, *Ichthyostega sp*.^2,20^. Ribs are depicted in purple. Pelvic girdles are shown in blue.

The rib anatomy of *Eusthenopteron*, coupled with a small ilium positioned ventrally to the vertebral column, indicate the absence of any linkage between axial column and pelvic fin^8,24^. The sacral ribs of *Tiktaalik*, on the other hand, would have overlapped the pelvic girdle in lateral perspective, with ribs lying medial to a large, plate-like ilium. Although there is no evidence of a bony articulation, the nature of the expansion of both ribs and ilium, the degree of overlap between the elements, and the unfinished distal margin of one sacral rib, indicates that a soft-tissue connection was likely in *Tiktaalik*. Such a connection, also proposed to be present in early limbed forms including *Acanthostega*^1^, likely allowed for a degree of structural support and for a restricted range of motion between the elements. A soft-tissue linkage between girdle and axial column would have provided a less robust a connection than direct bony articulations hypothesized for *Ichthyostega*^4^ and observed more clearly in more crownward forms, like *Whatcheeria*^19^. However, mobility of the pelvic girdle could have allowed for slight changes in the orientation of the acetabulum during locomotor behaviors. The post-cranial skeleton of *Tiktaalik*, therefore, reveals that sacro-iliac specializations arose in the ribs and pelvis prior to modifications to the vertebral column. Subsequent modifications to the axial column observed in limbed vertebrates include expansion of the dorsal extent of neural arches, either squared as in *Acanthostega* or rounded as in *Ichthyostega*, and the origin of zygapophyses^1,2^.

The presence of sacral ribs, robust pelvis, deep acetabulum, and large pelvic fin in *Tiktaalik* indicate that the rear appendage was generating greater forces in locomotion than in other finned elpistostegalians, such as *Panderichthys*. In addition, these features suggest that *Tiktaalik* was capable of more axial support for the trunk when the pelvic fins were loaded against the substrate than less crownward elpistostegalians. Despite these apomorphic features, *Tiktaalik* retains numerous plesiomorphic characteristics in its pelvic anatomy, such a posteriorly facing acetabulum, left and right pubes unfused along the midline, and lack of an ischium^14^, which imply that the pelvic fin was not able to retract as extensively as limbed forms such as *Acanthostega* and *Ichthyostega*. The posterior orientation of the acetabulum of *Tiktaalik* and concomitant inability to use retraction for limb propulsion suggests that the pelvic fin was unable to play a significant role in terrestrial walking.

With a pelvis and pelvic fin subequal in size to the shoulder girdle and pectoral fin, the overall proportions of the trunk and paired appendages of *Tiktaalik* hew closer to those of *Acanthostega*^1^ and *Ichthyostega*^2,20^ than to *Eusthenopteron*^8,24^ and *Panderichthyes*^11,15^. Pelvic and sacral anatomy implies that *Tiktaalik* represents an intermediate condition in which a large pelvic appendage was stabilized by the axial skeleton and capable of being used in diverse paddling, walking, and propping behaviors on aquatic substrates. These functions of the pelvic fin were antecedents to the terrestrial walking behaviors that were possible in later forms.

## Acknowledgments

Fieldwork was made possible by the Polar Continental Shelf Project of Natural Resources, Canada; Department of Heritage and Culture, Nunavut; the hamlets of Resolute Bay and Grise Fiord of Nunavut; the Iviq Hunters and Trappers of Grise Fiord. Douglas Stenton and Sylvie Leblanc (Department of Heritage and Culture, Nunavut) assisted with paleontology permits. Kieran Shepherd of the Canadian Museum of Nature assisted with export permits. We thank C F. Mullison for the original mechanical preparation of NUFV 108.

## Author contributions

Field work leaders: NHS, EBD; Funding acquisition: NHS, EBD; μCT data collection: JBL; μCT data processing: TAS; Visualization: TAS, EJH, IM; Writing – original draft: TAS, NHS; Writing – review & editing: TAS, JBL, EJH, IM, EBD, NHS

## Data and materials availability

All data analyzed in the paper will be freely available. Computed tomography data sets and STL files will be available for download from MorphoSource prior to publication.

## Supplementary Materials

### Materials and Methods

The material of *Tiktaalik roseae* was recovered during paleontological excavations near Bird Fiord on southern Ellesmere Island in 2004, 2006, 2008, and 2013. All specimens were recovered from a single locality (NV2K17; N77°09.895′ W86°16.157′) within the Fram Formation (Frasnian Stage, Late Devonian). The fossil material is curated in the Nunavut Fossil Vertebrate Collection (NUFV) at the Canadian Museum of Nature.

#### Computed tomography scanning

CT scans were collected at The University of Chicago’s PaleoCT scanning facility with a GE Phoenix vļtomeļx 240 kv/180 kv scanner. The post-cranial skeleton of NUFV 108 is contained in two blocks (Movie S1). Each of these blocks are too large for single multiscan. Therefore, each block was scanned each twice: first oriented vertically with the anterior edge down, and then rotated 180 degrees and scanned again with the posterior edge down. Scanning parameters for these four scans are provided in Table S1. CT data were reconstructed with Phoenix Datosļx 2 (version 2.3.3) and imported to VGStudio Max (version 2.2) to be cropped and exported as a tiff stack. For each block, the two multi-scans were manually stitched together and then manually segmented in Amira (version 20.2) (Thermo Fisher Scientific).

#### Surface scanning

The pelvis of NUFV 108 was previously physically prepared from the specimen and, therefore, is not included in the μCT scans. A 3D model of the pelvis was generated by surface scanning a cast of the pelvis using a FARO Design ScanArm 1.0 at a resolution of 40-75 μm.

#### Images and Animations

Volumetric images of the segmented μCT data were generated using Amira (Fig. 1, Fig. 4, Fig. S1). All other renderings of skeletal elements are surface models, which were generated by exporting segmentation label fields from Amira as surface files, or directly by surface scanning, and visualized in Blender (version 3.3.1). Movies were created by first exporting animations as tiff stacks from Amira or Blender and then using Adobe Premier (version 13.12) to combine and edit the images into movies.

## Supplementary Text

In specimen NUFV 108, individual skeletal elements are three-dimensionally preserved with minimal deformation, and the specimen has settled during preservation. Previous studies of *Tiktaalik* have presented reconstructions of the cranial and pectoral fin skeleton based on μCT data ^*21,23*^. In this study, we describe the axial skeleton of *Tiktaalik* and use μCT data to produce a three-dimensional model of specimen NUFV 108 that contains nearly all skeletal elements known for this taxon. Additional elements known from *Tiktaalik* that are not included in the model include components of the hyoid skeleton and the interclavicle. In generating our model, numerous decisions were made on how to place the pieces. These decisions are based upon comparisons between anatomical systems of *Tiktaalik* (*e.g*., comparing intercentral and neural arch anatomy, or comparing the pectoral and pelvic girdle), as well as comparisons to other tetrapodomorphs and extant fishes. The new reconstruction of *Tiktaalik*, thus, represents a hypothesis based on multiple lines of evidence.

### Reconstruction of the vertebrae

Intercentra and neural arches are positioned according to their preserved rostrocaudal order. Intercentra are assigned to either the left or right sides based curvature of the internal surface and position of the articular facet. Left and right intercentra that were preserved near to one another are reconstructed as paired. However, because intercentra are unfused can have shifted during preservation, it is possible that some elements reconstructed as paired are slightly out of register from their original position. Intercentra are reconstructed as associated with individual neural arches. Likewise, it is possible that intercentra could be reconstructed modestly out of register from their original neural arch. This uncertainty does not impact results presented in the manuscript.

Intercentra are positioned so that they bound the lower portion of the notochord and wrap dorsally. When both left and right sides are preserved for a vertebra, they are positioned so that their internal curvature symmetrically fits around a notochord that is circular in cross section. Intercentra are positioned under the assumption that the notochord is of a uniform cross section between the head and pelvis, a feature observed in various taxa, including *Eusthenopteron ^8,24^* and *Latimeria* ^29^. If only one intercentra was preserved for a vertebra, the element is positioned so its internal curvature matched elements anterior or posterior it in the series.

Neural arches are occasionally broken, and whenever possible the pieces are re-assembled. Neural arches from vertebrae 5-34 can be associated with ribs or intercentra. However, four neural arches are preserved more caudally and without clear association to other axial elements. The anterior-most of these four neural arches is preserved caudal and slightly ventral to neural arch 32 (Movie S1). Its morphology is significantly more robust than those immediately anterior (Fig. 2F-I), and it is identified as belonging to the caudal region based of comparison to *Acanthostega*^1^. Although the neural arch could have been associated with intercentra 34 or 35, it is depicted in the reconstruction with a gap between it and other elements to denote uncertainty in position (Fig. 2). The three most-caudal neural arches are preserved in close association with one another and separated by a substantial gap to other axial elements, near to the pelvic fin (Fig. 1A,B). These neural arches, too, are depicted in the reconstructed with gaps between them and other axial elements to denote ambiguity in their position (Fig. 2).

To reconstruct the dorsal position of neural arches, we first focused on the most complete neural arches in the series (*e.g*., Fig. 2F,G). Despite some lateral compression, these allowed us to estimate the extent to which the arch would have wrapped around the notochord. When neural arches were broken, if possible, they are reconstructed so that the apex of their internal curvature aligns with the apex of other more complete neural arches in the series.

To constrain spacing of axial elements in the rostrocaudal direction, we considered the preserved distance between ribs 1 and 32 in NUFV 108 to approximate the distance between ribs 1 and 32 in life. Across this distance, ribs and vertebrae are placed so that gaps between the vertebrae were uniform, except when their position was uncertain (see discussion above of the caudal-most 4 vertebrae and discussion below of the sacral rib). Vertebrae 32-36 are spaced at distances similar to those of positions 1-32.

In the reconstruction of *Tiktaalik*, intercentra are positioned slightly anterior to their corresponding neural arch. This positioning is based on several features. First, the positioning of intercentra reveals the size of the notochord, and comparison of intercentral and neural arch morphologies suggest that they are unlikely to have been aligned strictly dorsally, because this would have produced a lateral overlap of the elements. Second, pleurocentra were not identified for specimen NUFV 108. If large pleurocentra had been observed in the specimen, then the vertebrae are likely to have been organized such that neural arches were positioned dorsal to their corresponding intercentra, as in *Osteolepis* ^9^. Therefore, the absence of pleurocentra indicates they were either small or fused to the intercentra; both conditions predict that neural arches and intercentra were not vertically aligned, but slightly out of register ^3,8^.

The neural arches of *Tiktaalik* lack zygopophyses. This suggests space between adjacent neural arches. Therefore, they are situated with angles of inclination that maintain a slight gap between adjacent elements. The caudal four neural arches are positioned with similar angles of inclination as those in the trunk series.

### Reconstruction of the ribs

The anterior-most rib on the left side is broken in two pieces, which were preserved in contact with one another with a sharp angle between them (Fig. 1). These pieces are placed end-to-end to reconstruct the original element (Fig. 2). Other ribs that are broken have pieces preserved in close proximity with one another, and they are approximately aligned (*e.g*., rib 23 on the right side). In the reconstruction, the pieces of these other broken ribs are kept in their preserved positions and have not been moved closer to one together in the reconstruction. This presentation was done to preserve information on which features are broken and not to imply that any gaps in individual ribs represent their original length and missing portions of the rib.

Two ribs on the left side (ribs seven and twelve) and one on the right side (rib six) were displaced during preservation such that the distal portion of the rib was posteriorly oriented and ventral to the rib that followed. Additionally, four ribs on the right side (ribs 20-23) are preserved such their articular surfaces point posteriorly. In each of these cases, the individual ribs were rotated and repositioned preserving the order of their proximal articular surfaces.

One rib is preserved to the left of the rest of the axial series, and it is identified as a post-sacral rib. It is possible that it might have articulated upon intercentra 33-36, as approximately 5 post-sacral ribs are preserved in *Acanthostega ^1^* and *Ichthyostega ^2,20^*. However, the rib is depicted in the reconstruction with a gap between it and other axial elements to denote ambiguity in position.

Ribs were positioned relative to the vertebral column based on curvature of the proximal articular surface. In many ribs, this portion is broken or incomplete. Therefore, across the series, ribs are placed by first reconstructing the positions of those ribs with complete articular heads. These ribs were placed so that their heads aligned with the curvature of the posterior margin of the intercentra, which bear an articular facet. Ribs with damaged heads were then positioned to maximize similarity in their orientation to those with complete heads.

### Reconstruction of the pelvis

Rostrocaudal positioning of the pelvis of *Tiktaalik* is based upon transitions in rib and neural arch anatomy. Specifically, the girdle is placed so that the dorsal extent of the ilium is rostrocaudally aligned with the sacral ribs (ribs 31 and 32). This positioning is similar to what has is proposed for the pelvic girdles of *Acanthostega* and *Ichthyostega*, which each have approximately 30 pre-sacral vertebrae ^1,2^.

Dorsoventral positioning of the pelvic girdle of *Tiktaalik* is based on comparisons to other tetrapodomorphs. Uniformly, tetrapodomorphs are reconstructed with the ventral portion of the pelvic girdle approximately in line with the ventral portion of the pectoral girdle (*e.g.*, *Eusthenopteron* ^8^, *Acanthostega* ^1^, *Ichthyostega* ^2^). In *Tiktaalik*, thus, the pelvic girdle is placed with a position that comports with the body thickness observed in the articulated pectoral region.

To reconstruct the medio-lateral splay of the pelvic girdle of *Tiktaalik*, first the anteromedial portion is positioned near to the midline, as in *Eusthenopteron* ^8^. Next, the girdle was positioned to produce a taper in the body outline when viewed from the dorsal perspective. Specimen MHNM 06-2067 of *Elpistostege* shows approximately 30% reduction in the width of the trunk between the pectoral and pelvic fins ^30^. The pelvic girdle of *Tiktaalik* is reconstructed similarly, resulting in a narrow distance between the ilium and sacral ribs (Fig. S1 A-E). This reconstruction predicts a more posterior orientation of the acetabulum than previously hypothesized ^14^, one approximately similar to *Eusthenopteron* ^8^. We regard this hypothesis of pelvic positioning as more likely than one where the dorsal extent of the ilium is parallel to the axial column (Fig. S1 F-I). Such a wide splay would result in an unusually ovate shape of the trunk in cross section at the position of the pelvis (Fig. S1 H), which is not known among tetrapodomorphs. Additionally, if a lateral orientation is constrained, but the angle between left and right halves is increased to produce a more rounded cross-section, this increases the height of the girdle in lateral perspective and yields a reconstruction where body thickness is greater at the pelvis than the pectoral girdle (Fig. S1 H). As noted above, such an increase in body thickness is not seen in other closely related taxa and regarded as unlikely.

Thus, positioning of the pelvis is constrained by both features of other anatomical systems (*i.e*., vertebrae, ribs, and pectoral girdle) and by comparisons to other tetrapodomorphs. Although there is uncertainty in some features of the reconstruction, alternative hypotheses of pelvic girdle positioning for *Tiktaalik* robustly predict that the dorsal extent of the ilium approached the sacral ribs and that they overlapped in lateral perspective. Further, alternative predictions also recover a pelvic fin of *Tiktaalik* that is more posteriorly oriented than in *Acanthostega* and *Ichthyostega*^1,2^.

### Reconstruction of the pelvic fin

A line drawing of the pelvic fin is presented in Fig. 4 A that shows the estimated positions of the preserved endoskeleton elements as well as estimates of the geometry of missing elements. Along the proximodistal axis, fins generally taper dorsoventrally. Accordingly, proximal skeletal elements have articular surfaces that are deeper in the dorsoventral direction than those more distally positioned. As previously noted, element shown in purple in Fig. 4 has a similar morphology to the intermedium of the pectoral fin of *Tiktaalik*^14^; it is, thus, reconstructed as articulating with the fibula. This positioning contributed to the identification of the tibia. The element identified as the tibia has an articular surface deeper dorsoventrally than any other preserved pelvic endoskeletal elements and, therefore, would likely have been more proximally positioned than the element shown in purple. The general pattern of tetrapodomorph pelvic fins is such that one would predict only three possible more proximal elements: the femur, fibula, and tibia. The geometry of this most robust element is inconsistent with either a femur or fibula, both of which likely would have had two distal articular facets, and it is therefore identified as the tibia.

In the drawing, the tibia is illustrated with a dashed component distal to it. The distal geometry of the tibia is rough and uneven as compared to the distal surfaces of other pelvic elements, like the intermedium, third mesomere, and third anterior radial. Therefore, this texture is taken to indicate that the distal portion of the tibia might have broken off or was poorly ossified. It is possible that a small element articulated distally with the tibia. We regard this condition as unlikely, because neither *Eusthenopteron* ^8^ nor *Panderichthys* ^15^ have pelvic fins showing an element articulating distally with the tibia.

Several endoskeletal elements of the pelvic fin are not preserved. Their approximate geometries are estimated in the illustration. Mesomeres are typically not longer proximodistally than those more proximal to them. Therefore, we estimated the relative lengths of the fibulare, fibula based on the third mesomere (shown in red in Fig. 3). The approximate geometry of the femur is based on the assumptions that it would be at least as long as the tibia and distally wide enough to accommodate the tibia and fibula.

In the pectoral fin, fin rays overlap with the radius. The tibia, the homologous element in the pelvic fin, is therefore expected to similarly have been covered by lepidotrichia. Accordingly, it is positioned it in the 3D reconstruction so that dorsal hemitrichia would have reached approximately to the base of the femur. Individual fin rays within the fin web are not repositioned. The pelvic fin is placed relative to the girdle such that a femur, if present, would be extending straight from the acetabulum.

---

**Fig. S1.**
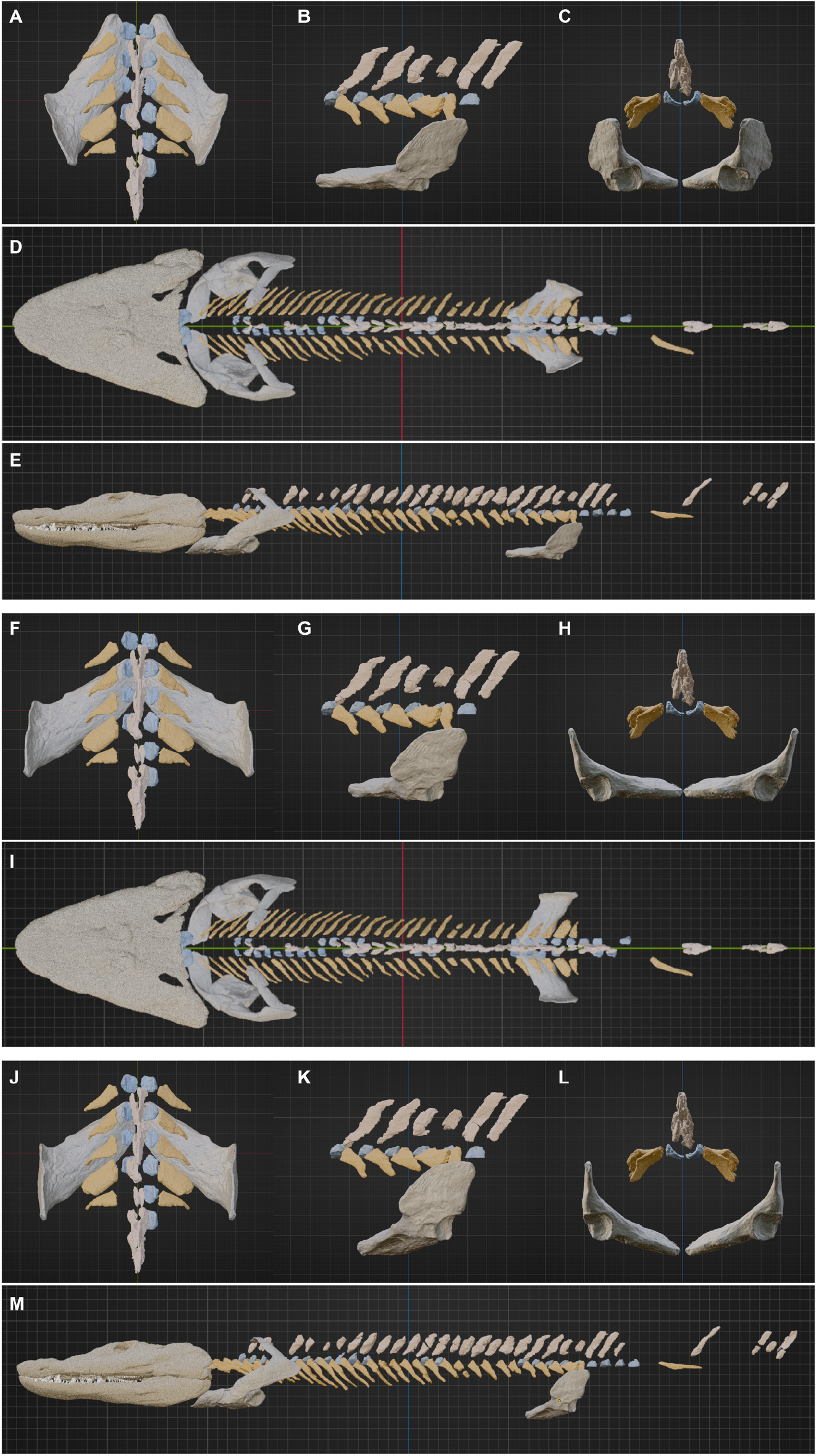
Alternative reconstructions of pelvic girdle of *Tiktaalik roseae*. Alternative hypotheses for the positioning of the pelvic girdle were considered when building the reconstruction, as reviewed in the Supplementary Discussion. Panels A-E show the reconstruction of the pelvic girdle presented in the main manuscript. Panels F-I compare that condition with an alternative positioning, where the dorsal extent of the ilium is parallel to the rostro-caudal axis and the ventral aspect of the pelvic girdle is aligned with the ventral aspect of the pectoral girdle. This position, with a broad body in the pelvic region, corresponds to previous reconstruction of the pelvic girdle^14^. Panels J-M show a third reconstruction, where the left and right halves of the pelvis are rotated.

**Fig. S2.**
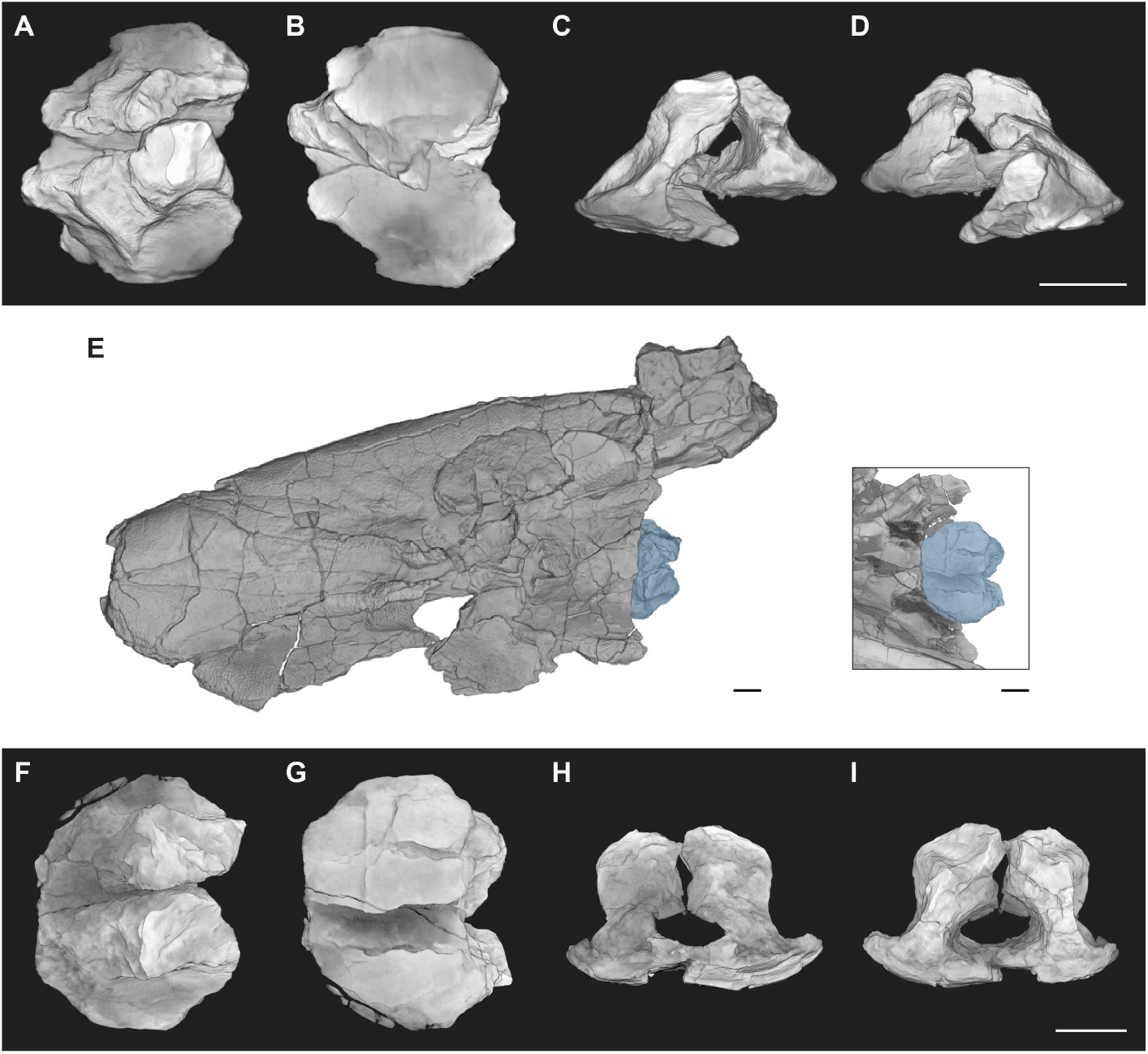
Basioccipital-exoccipital complex of *Tiktaalik roseae*. The basioccipital-exoccipital complex of *Tiktaalik* is preserved in specimens NUFV 108 and NUFV 110 as paired elements that are unfused to the rest of the braincase. In NUFV 108, the elements are preserved medial to the pectoral girdle, as depicted in Fig. 1. The basioccipital-exoccipital elements of NUFV 108 shown in preserved positions from (A) dorsal, (B) ventral, (C) anterior, and (D) posterior perspectives. (E) In NUFV 110, the basioccipital-exoccipital complex is still contacting the rest of the skull. The basioccipital-exoccipital complex of NUFV 110 from (F) dorsal, (G) ventral, (H) anterior, and (I) posterior perspectives. Scale bars, 1 cm.

**Fig. S3.**
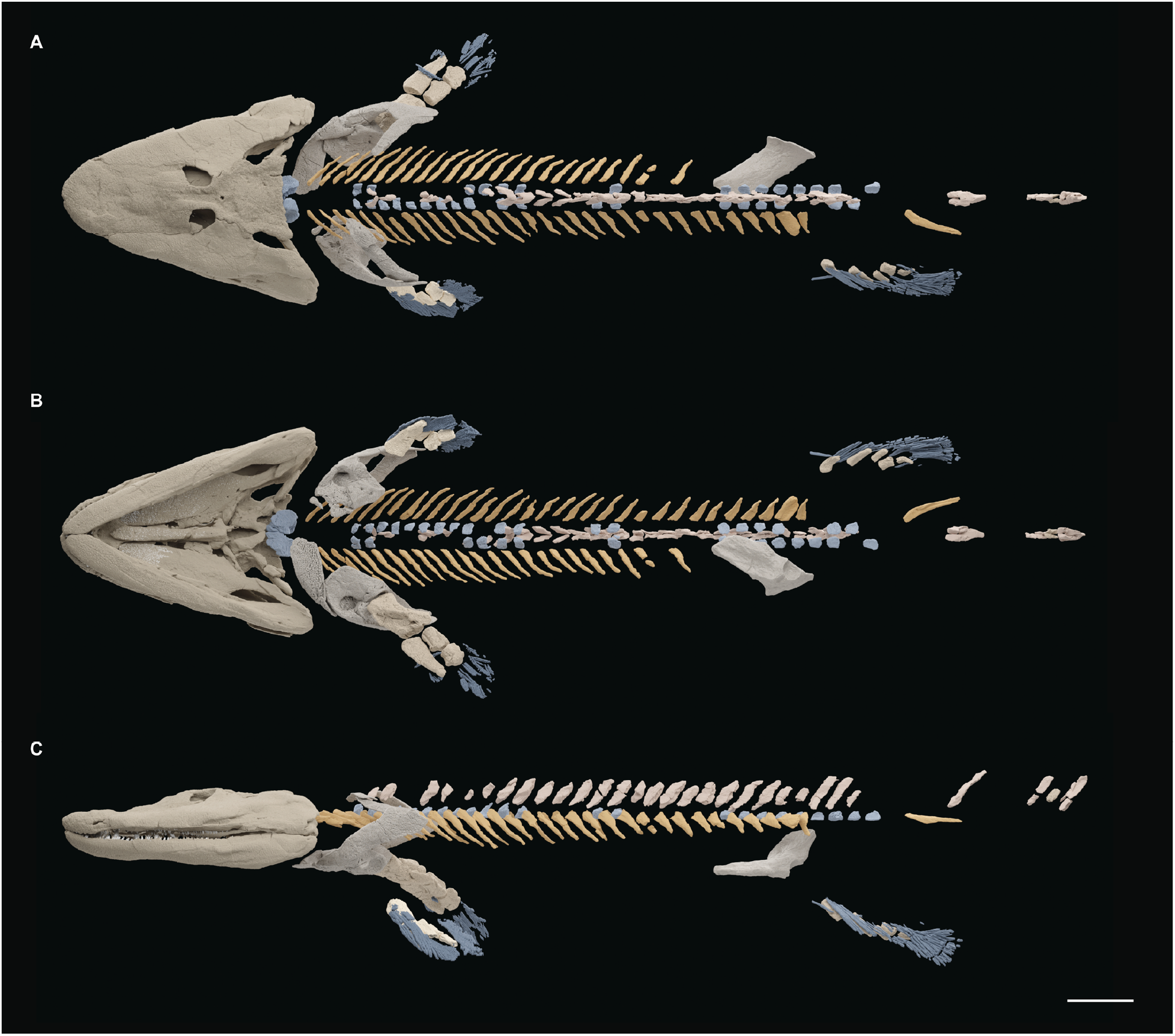
Specimen NUFV 108 with elements repositioned. Rendering of all skeletal elements of NUFV 108 that have been μCT scanned are shown here in their reconstructed positions. These images differ from the reconstruction in Fig. 5, which shows several elements duplicated for left-right symmetry and coupled with the more complete pectoral fin of another specimen. Scale bar, 5 cm.

**Table S1.** Parameters for μCT scanning of specimen NUFV108

| scan ID | Voltage (kV) | Current (μA) | Voxel Size (μm) | Filter (mm) | Scan Duration |
| --- | --- | --- | --- | --- | --- |
| pectoral<br>block<br>anterior | 100 | 570 | 122.441 | 0.24 Cu | 1hr44min |
| pectoral<br>block<br>posterior | 100 | 570 | 88.038 | 0.24 Cu | 5hr06min |
| pelvic<br>block<br>anterior | 110 | 450 | 73.23 | 0.24 Cu | 6hr48min |
| pelvic<br>block<br>posterior | 110 | 450 | 73.23 | 0.24 Cu | 6hr48min |

Parameters for μCT scanning of specimen NUFV108

**Movie S1.**

Volumetric rendering of the two blocks containing the post-cranial skeleton of NUFV 108 including matrix

**Movie S2.**

Volumetric rendering of NUFV 108 with all segmented elements in their preserved position

**Movie S3.**

Rotation of the reconstructed sacral domain of *Tiktaalik roseae*

**Movie S4.**

Rotation of the reconstruction of *Tiktaalik roseae*

